## Supplementary Text for "A deep population reference panel of tandem repeat variation"

### Supplementary Methods

#### Experimental validation of TR genotypes

The following loci were amplified using separate PCR conditions than those described in **Methods**:

- SCA1(ATXN1): 95°C for 5 mins, 28 cycles of 95°C for 1 min, 61°C for 1 min, 72°C for 1.5 min; 8 cycles of 95°C for 1 min, 53°C for 1 min, 72° C for 1.5 mins, and a final extension at 72°C for 10 min.
- ATN1: 95°C for 15 min, 27 cycles at 94°C for 20 s, 65° C for 30 s, 72°C for 1.5 min; 8 cycles of 94°C for 20 s, 53°C for 30 s, 72°C for 1.5 mins and a final extension at 72°C for 10 min.

Several additional loci were initially analyzed by capillary electrophoresis but were filtered from downstream analysis since we determined they produced unreliable genotypes:

- Known pathogenic repeats at *HOXD13* (F:tgtaaaacgacggccagtAAGGAGAGGAGGGAGGAGG GACATACGGCAGCTGTAGTAGC, R:GACATACGGCAGCTGTAGTAGC and *PAPBN1* (F:tgtaaaacgacggccagtCCAGTGCCCCGCCTTAGA, R: ACAAGATGGCGCCGCCGCCCCGGC) were excluded since all samples were called as homozygous reference by both EnsembleTR and capillary electrophoresis.
- A known pathogenic repeat in FXN was genotyped but excluded from comparisons since it was not genotyped by EnsembleTR.
- A known pathogenic repeat for SCA2 (F:6FAM-GGGCCCCTCACCATGTCG, R:CGGGCTTGCGGACATTGG) was excluded since many capillary calls had evidence of three alleles in a single sample, and we could not determine the correct diploid call.
- A known pathogenic repeat in *PHOX2B* (F:tgtaaaacgacggccagtCCAGGTCCCAATCCCAAC R:GAGCCCAGCCTTGTCCAG) was excluded since all capillary calls were heterozygous for the same alleles, which is unlikely to be correct.
- We additionally excluded known pathogenic loci on chrX, since our calls focus on autosomal TRs. Results for the FMR1 repeat, which is on chrX, are reported from Asuragen but are not included in comparison.

Fragment analysis was performed in separate batches for Platinum Genomes (17 samples; "PG") and the remaining 31 samples ("Coriell"). Product sizes were converted to repeat differences from reference to be comparable to our WGS genotypes. In some cases, product sizes were offset by a constant value from the expected values. We binned product sizes to allele lengths, with binsets manually set for each locus in each batch. For each locus, we set the minimum (minval) and maximum (maxval) product size, the repeat unit length (period), and allele assignment of the minimum product size (start). We then determined the allele for a given product size (psize) using the following rule:

1. Set allele=start and i=minval
2. While psize >= i and psize < i+period and i <= maxval: allele += 1
3. If psize >= i and psize < i+period: return allele. Else, return NA

The bin definitions for each locus/batch are provided in **Supplementary Table 5** below.

**Supplementary Table 5: Bin sets used to convert PCR product sizes to allele lengths.**

| **Locus** | **Cohort** | **Minval** | **Maxval** | **Period** | **Start** |
|---|---|---|---|---|---|
| ATN1 | PG+Coriell | 129 | 190 | 3 | -7 |
| ATXN10 | PG+Coriell | 192 | 240 | 5 | -3 |
| CACNA1A | PG+Coriell | 131 | 170 | 3 | -7 |
| chr1_106950468_GT* | PG+Coriell | 177<br>197 | 197<br>220 | 2<br>2 | -6<br>3 |
| chr1_217937145_AAAT | PG+Coriell | 174 | 200 | 4 | -5 |
| chr1_242233155_AATG | PG+Coriell | 224 | 300 | 4 | -1 |
| chr1_6733191_CA | PG+Coriell | 168 | 200 | 2 | -1 |
| chr1_76108862_GT | PG+Coriell | 244 | 300 | 2 | -11 |
| chr10_57337770_CA | PG+Coriell | 196 | 250 | 2 | -7 |
| chr10_72427734_AT | PG+Coriell | 198 | 250 | 2 | -10 |
| chr11_2442377_AC | PG+Coriell | 171 | 250 | 2 | -5 |
| chr11_36246191_AATA | PG+Coriell | 191 | 250 | 4 | -3 |
| chr11_63798227_AC | PG+Coriell | 234 | 300 | 2 | -5 |
| chr11_85908552_TAGA | PG+Coriell | 204 | 300 | 4 | -5 |
| chr12_70011632_AG | PG | 225 | 260 | 2 | -4 |

| | | | | | |
|---|---|---|---|---|---|
| | Coriell | 226 | 260 | 2 | -4 |
| chr13_102648430_GT | PG+Coriell | 214 | 250 | 2 | -7 |
| chr13_75404064_AC | PG+Coriell | 228 | 280 | 2 | -4 |
| chr13_81527591_CTAT | PG+Coriell | 296 | 350 | 4 | -1 |
| chr15_53480481_AC | PG+Coriell | 299 | 350 | 2 | -7 |
| chr15_80839290_GT | PG+Coriell | 203 | 250 | 2 | 0 |
| chr16_62573638_AATA | PG+Coriell | 249 | 300 | 4 | -2 |
| chr17_8128309_TAGA | PG+Coriell | 380 | 420 | 4 | -3 |
| chr18_38020886_TCTA | PG+Coriell | 380 | 450 | 4 | -5 |
| chr2_1273636_TA | PG+Coriell | 234 | 280 | 2 | -1 |
| chr20_18095896_AC | PG+Coriell | 178 | 250 | 2 | -9 |
| chr20_42508128_TG | PG+Coriell | 233 | 280 | 2 | 0 |
| chr3_46654755_TG | PG+Coriell | 312 | 350 | 2 | -2 |
| chr4_123573491_TTAT | PG+Coriell | 201 | 250 | 4 | -1 |
| chr4_136965932_AT | PG+Coriell | 176 | 250 | 2 | -3 |
| chr4_46950717_AAAT | PG+Coriell | 304 | 350 | 4 | 0 |
| *chr5_18768193_GT | PG+Coriell | 157<br>176 | 177<br>250 | 2<br>2 | -8<br>1 |
| chr6_12322154_AC | PG+Coriell | 258 | 300 | 2 | -3 |
| chr6_50207587_TCTA | PG+Coriell | 400 | 450 | 4 | -2 |
| chr6_84874972_AC | PG+Coriell | 239 | 300 | 2 | -2 |
| chr6_89959098_CA | PG+Coriell | 210 | 300 | 2 | -11 |

| | | | | | |
|---|---|---|---|---|---|
| chr7_18349308_AC | PG<br>Coriell | 206<br>207 | 300<br>300 | 2<br>2 | -7<br>-7 |
| chr7_27264534_AC | PG+Coriell | 224 | 250 | 2 | -4 |
| chr8_50595021_TG | PG+Coriell | 224 | 275 | 2 | -3 |
| chr9_36061394_AC | PG+Coriell | 250 | 300 | 2 | -6 |
| chr9_6685998_TTA | PG+Coriell | 265 | 300 | 3 | 0 |
| DMPK | PG<br>Coriell | 856<br>855 | 950<br>950 | 3<br>3 | -15<br>-17 |
| JPH3 | PG+Coriell | 113 | 200 | 3 | -6 |
| PPP2R2B | PG<br>Coriell | 162<br>159 | 200<br>200 | 3<br>3 | 0<br>-2 |
| SCA1 | PG+Coriell | 211 | 260 | 3 | -7 |
| SCA3 | PG+Coriell | 114 | 200 | 3 | 0 |
| SCA7 | PG<br>Coriell | 291<br>290 | 350<br>350 | 3<br>3 | 0<br>-1 |

* For locus chr1_106950468_GT, a separate offset was observed for alleles with product sizes <197bp. For locus chr5_18768193_GT, a separate offset was observed for alleles with product sizes < 177bp.

### Supplementary Figures

#### Supplementary Fig. 1

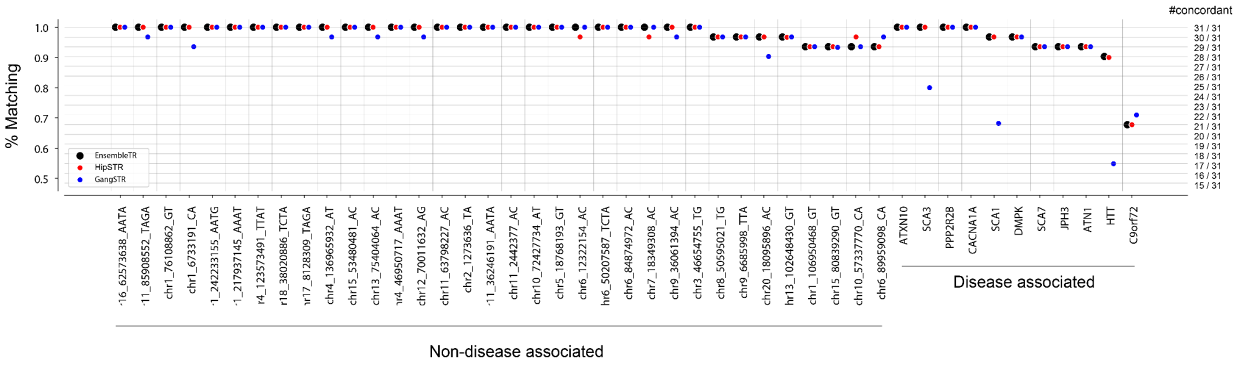


**Per-locus comparison between WGS-based TR genotypes and capillary electrophoresis results.** Each point gives the percent of genotypes that were concordant (y-axis) between capillary electrophoresis and WGS (black=EnsembleTR; red=HipSTR; blue=GangSTR) for each TR (x-axis). TRs are arranged by category (left=non-disease associated TRs; right=disease-associated TRs). Within each category, TRs are sorted by EnsembleTR concordance. Horizontal lines denote the % matching corresponding to the different possible numbers of concordant genotypes (out of a maximum of 31 samples). Full genotype comparison results are provided in **Supplementary Table 4**. Note, for HTT, GangSTR does not distinguish between the $CAG_n$ and $CCG_n$ repeats, whereas the Asuragen test used for PCR counts only CAG copies. Similarly, SCA1 consists of an imperfect repeat, and GangSTR only considers the length of perfect repeat stretches. This likely accounts for lower concordance of GangSTR compared to EnsembleTR and HipSTR at those loci (**Methods**).

Supplementary Fig. 2

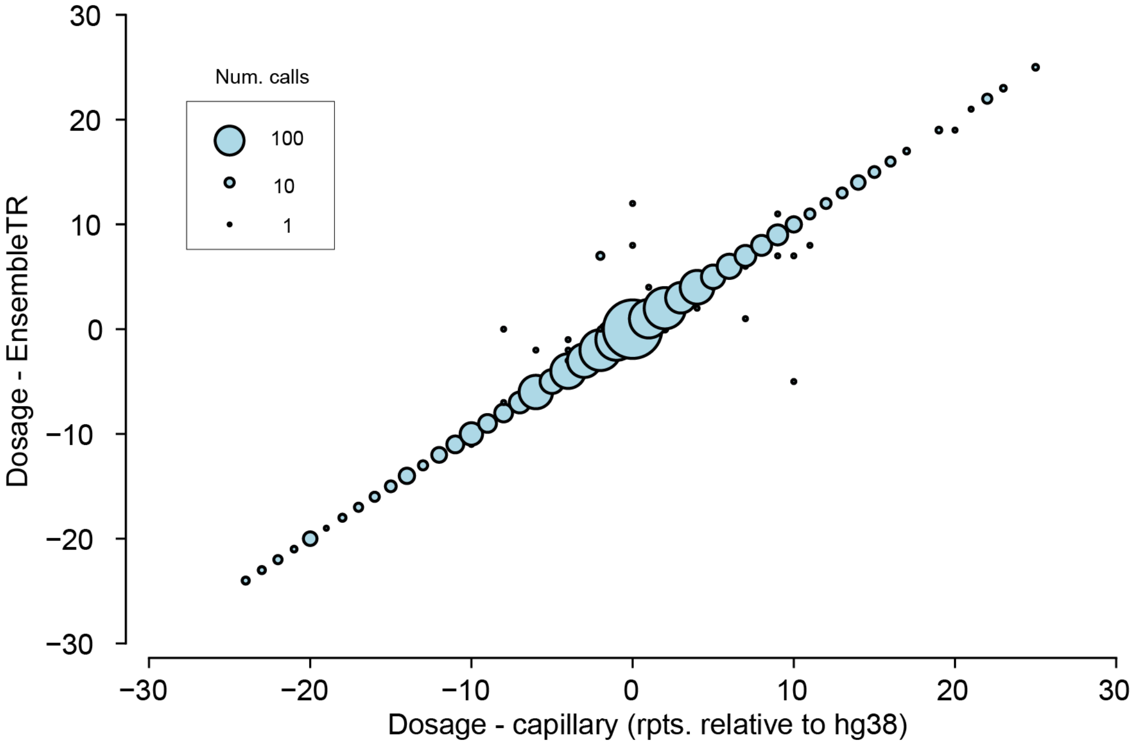


**Comparison of genotypes called by capillary electrophoresis vs. EnsembleTR.** The length of each allele relative to the hg38 reference was computed and lengths of the two alleles were summed for each call. The x-axis gives the capillary result, and the y-axis gives the result called by EnsembleTR. Bubble size scales with the number of genotype calls at each coordinate.

#### Supplementary Fig. 3

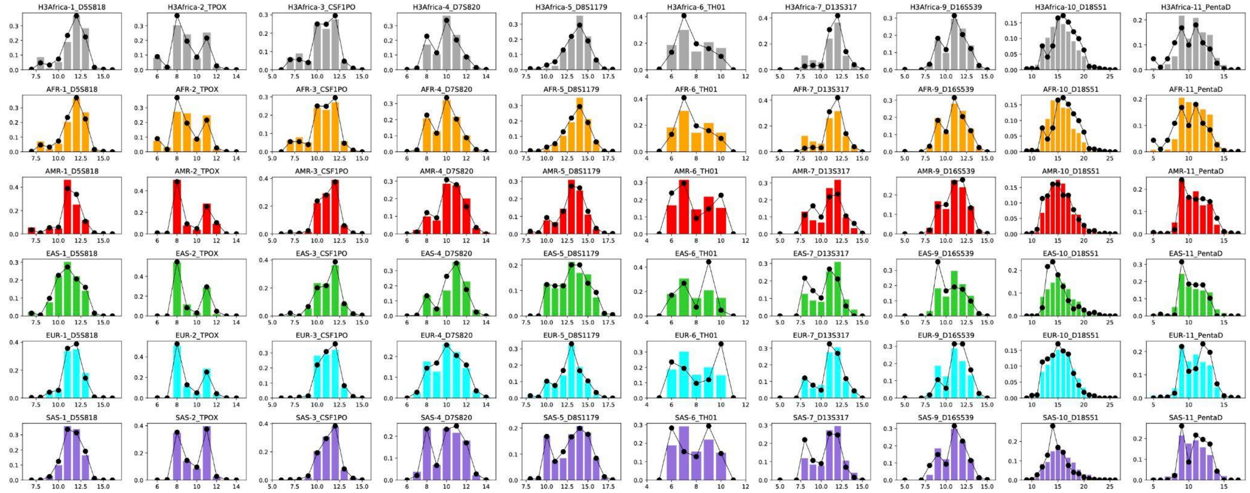


**Comparison of population-specific allele length distributions called by EnsembleTR vs. published frequencies at the CODIS forensic loci.** Rows represent 1000GP superpopulations and H3Africa (gray=H3Africa, blue=EUR, green=EAS, orange=AFR, red=AMR, purple=SAS). Each column represents a different CODIS locus. Colored bars give the frequency (y-axis) of each allele length (x-axis) observed in EnsembleTR results in terms of the number of repeat units. EnsembleTR alleles at the TH01 locus, a tetranucleotide repeat with a common 3bp interruption, were rounded down to the nearest repeat unit. All EnsembleTR alleles at D5S818 were adjusted down by one repeat unit since the EnsembleTR call contains an extra imperfect repeat unit not counted in published frequency data. Black dots give the expected frequency of each length based on previous literature (see **Methods**).

Supplementary Fig. 4

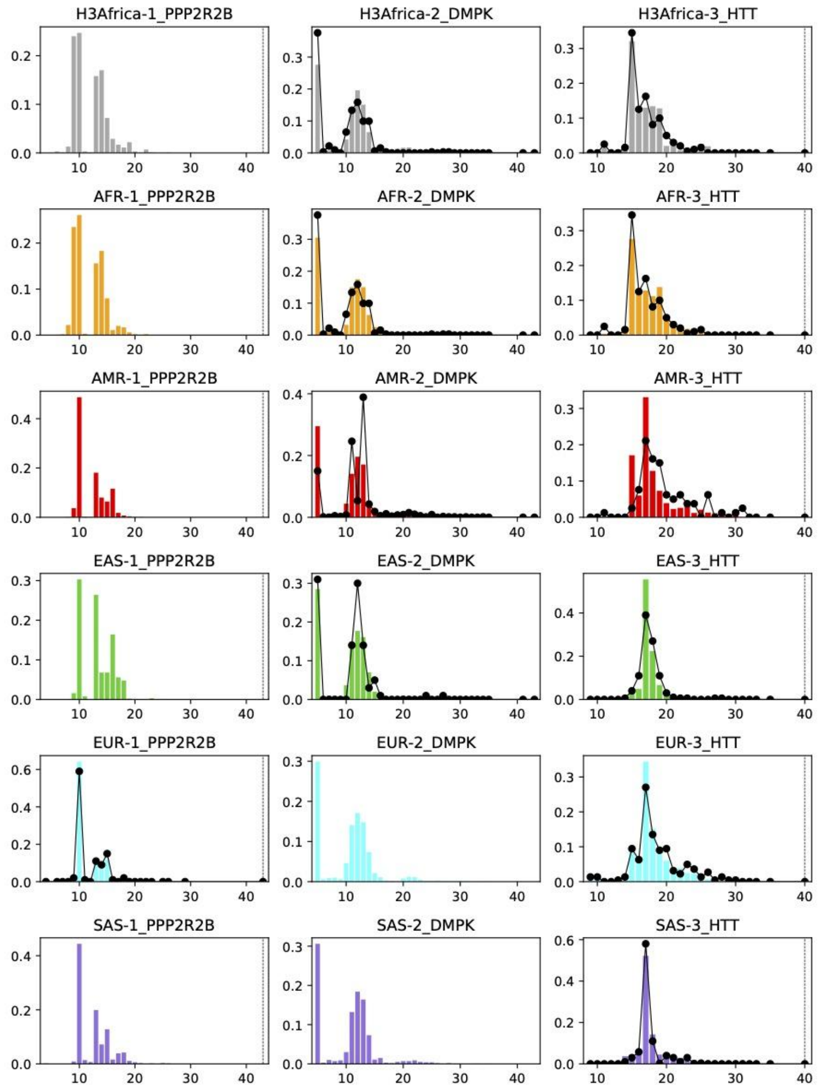


**Comparison of population-specific allele length distributions called by EnsembleTR vs. published frequencies at the disease-associated loci.** Rows represent 1000GP superpopulations and H3Africa (gray=H3Africa, blue=EUR, green=EAS, orange=AFR, red=AMR, purple=SAS). Each column represents a different locus implicated in a TR expansion disorder. Colored bars give the frequency (y-axis) of each allele length (x-axis) observed in EnsembleTR results in terms of the number of repeat units. Black dots give the expected frequency of each length based on previous literature (see **Methods**). For HTT, we counted only the number of CAG repeats in alleles returned by EnsembleTR to be consistent with counts obtained in the literature, which do not count the adjacent CCG repeat.

Supplementary Fig. 5

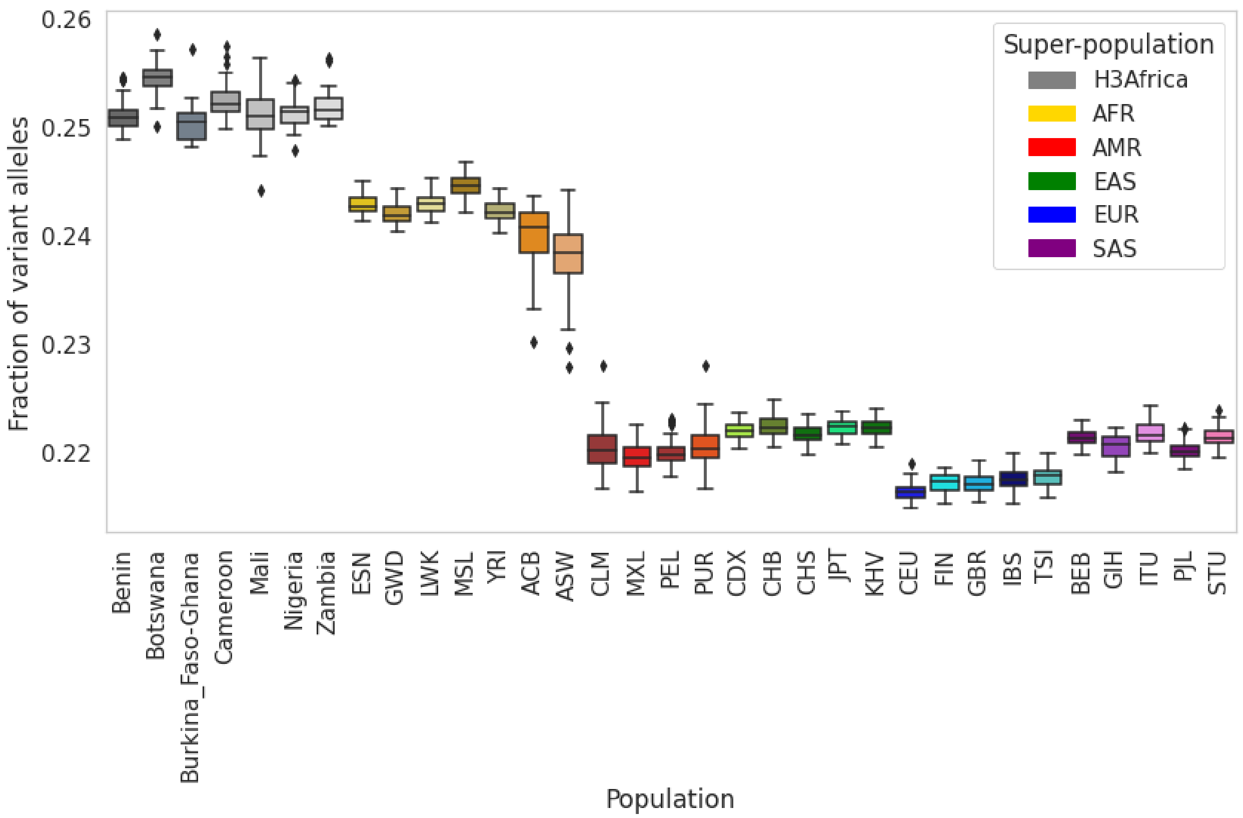


**Distribution of the fraction of non-reference alleles in individuals by population.** Boxplots summarize the distribution of the fraction of variant alleles in each sample. Horizontal lines show median values, boxes span from the 25th percentile (Q1) to the 75th percentile (Q3). Whiskers extend to Q1–1.5*IQR (bottom) and Q3+1.5*IQR (top), where IQR gives the interquartile range (Q3-Q1). Data is the same as in **Fig. 1d** except homopolymer TRs are included.

Supplementary Fig. 6

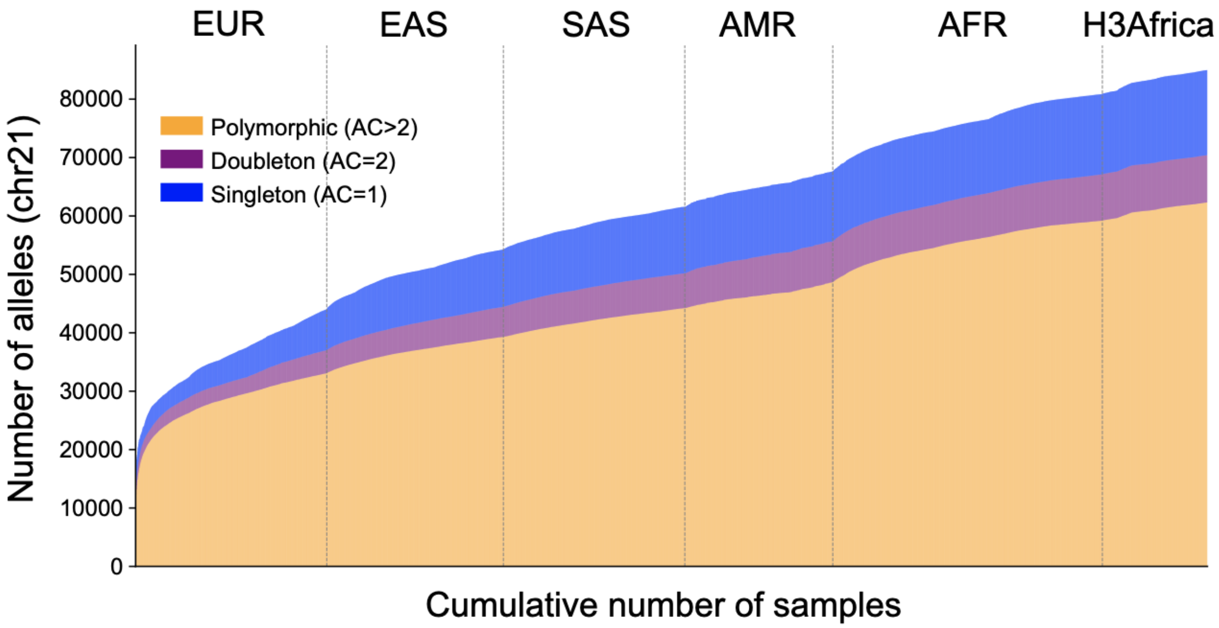


**Cumulative number of TR alleles with each additional sample.** The y-axis gives the number of unique TR alleles discovered with each additional sample. Samples on the x-axis are sorted by superpopulation. Orange=polymorphic alleles (allele count > 2), purple=doubleton (observed twice), blue=singleton (observed once). Data is based on chromosome 21.

#### Supplementary Fig. 7

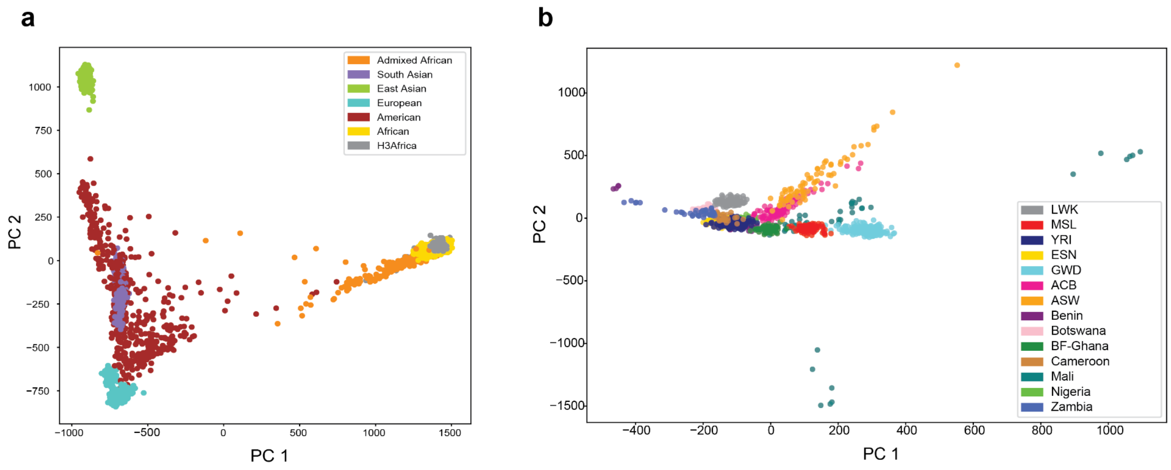


**TR variants capture known population structure. a. Principal components analysis of all samples.** PCA was performed using TR genotypes on all samples (**Methods**). The projection of each sample along PC1 and PC2 are shown. **b. Principal components analysis of African populations.** A separate PCA was performed on only the H3Africa and 1000GP AFR samples. Samples of similar origin cluster despite being sequenced and genotyped separately (e.g. YRI [Yorubans from Nigeria] from 1000G and Nigeria from H3Africa are clustered together). For **a-b**, each dot represents a single individual. Dots are colored based on the population or superpopulation of origin.

Supplementary Fig. 8

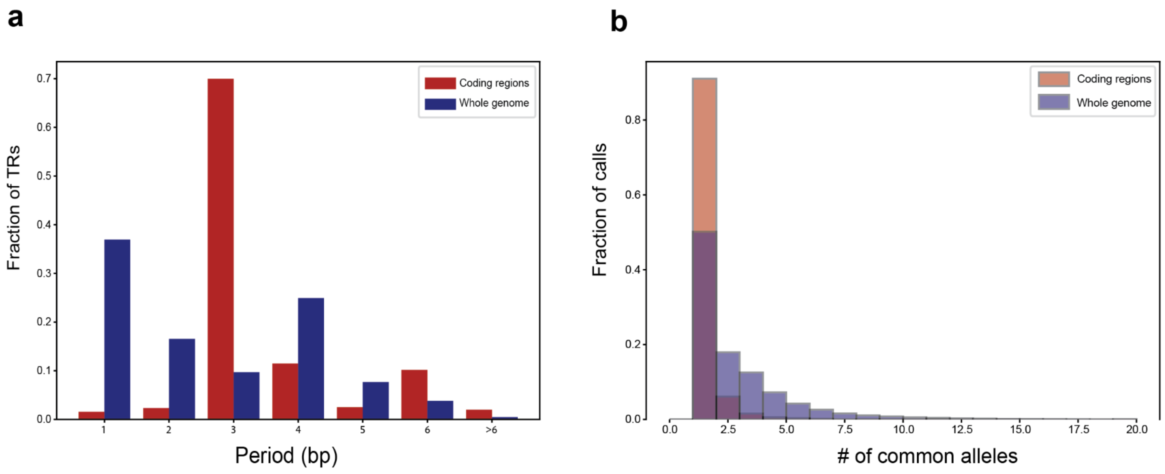


**Comparison of TR patterns in coding regions vs. genome-wide. a. Comparison of TR repeat unit length distributions.** While homopolymer and dinucleotide repeats are most prevalent genome-wide, trinucleotide TRs are over-represented in coding regions. **b. Comparison of the distribution of the number of common alleles at each locus.** Common alleles are defined as having a frequency > 1%. As expected, coding TRs are less polymorphic and have fewer common alleles per TR compared to genome-wide TRs. In both panels, red bars denote TRs in coding regions and blue bars denote genome-wide TRs.

#### Supplementary Fig. 9

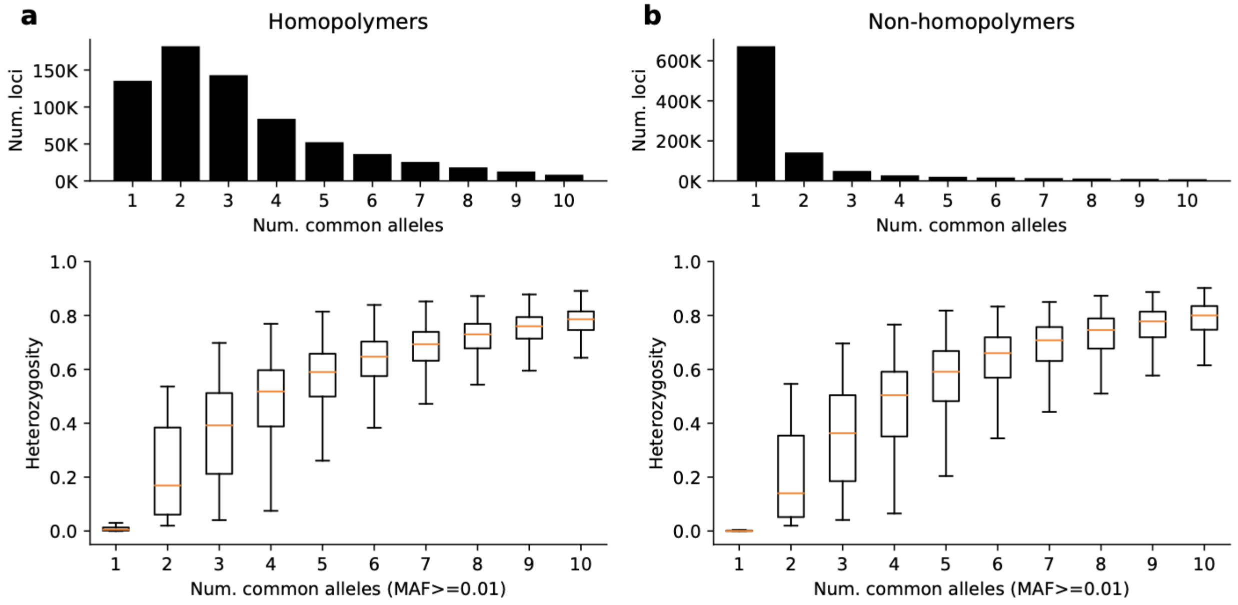


**TRs exhibit a range of polymorphism levels across loci.** **a.** shows data for only homopolymer TRs, and **b.** shows data for non-homopolymer TRs. Both panels show heterozygosity as a function of the number of common alleles. The y-axis gives the distribution of heterozygosity values for TRs with the number of common alleles specified in the x-axis. Boxplots summarize the distribution of heterozygosity values. Horizontal lines show median values, boxes span from the 25th percentile (Q1) to the 75th percentile (Q3). Whiskers extend to Q1–1.5*IQR (bottom) and Q3+1.5*IQR (top), where IQR gives the interquartile range (Q3-Q1). In **a-b**, top panels show the number of TRs with each number of common alleles.

Supplementary Fig. 10

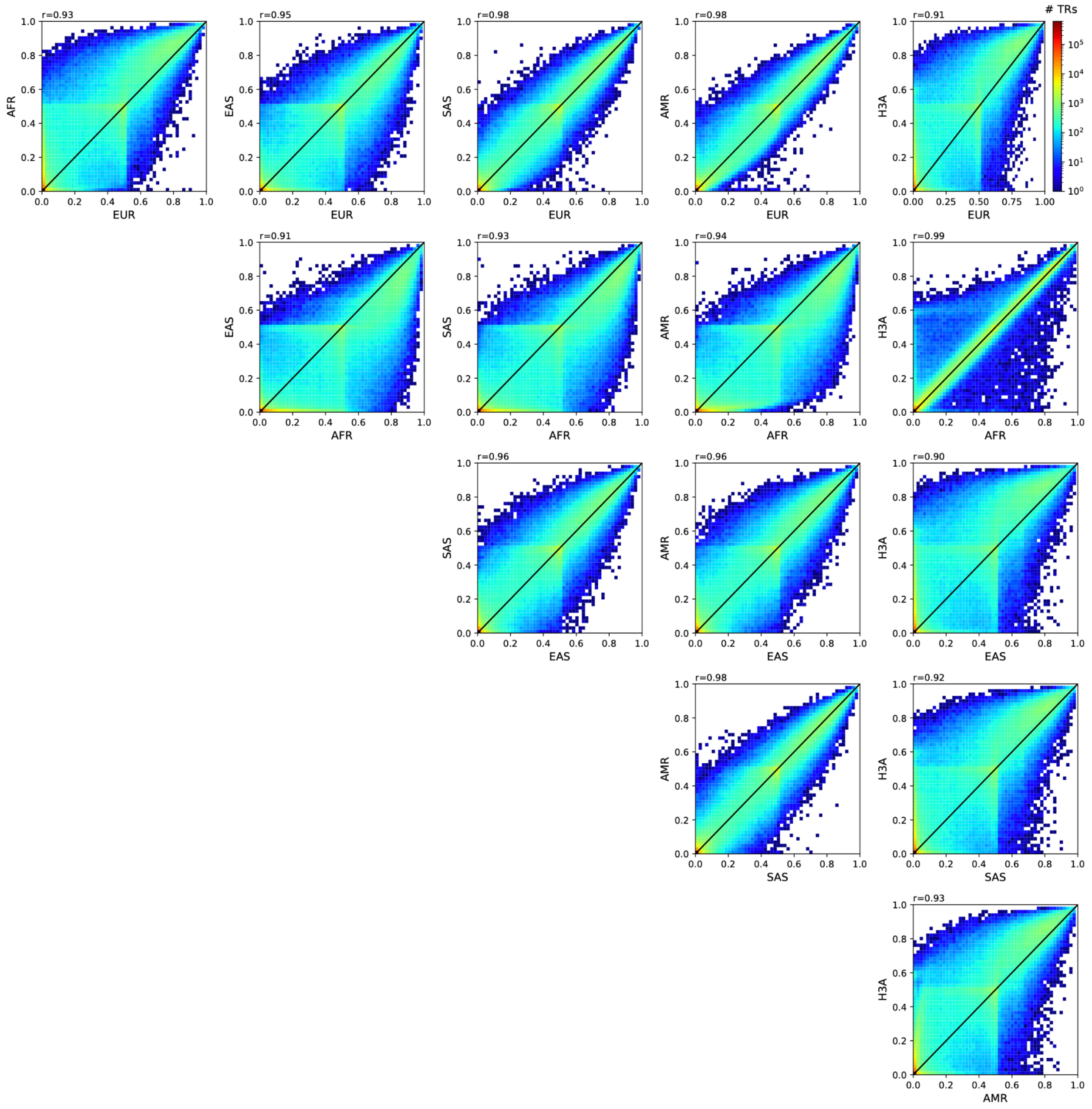


**Comparison of TR heterozygosity values across populations.** Each scatterplot compares the heterozygosity of each TR across two superpopulations. The black line denotes the diagonal. Heatmap shading indicates the number of TRs falling into each bin. The Pearson correlation comparing heterozygosity values in each pair of superpopulations is annotated above each panel. Data is shown for non-homopolymer TRs.

Supplementary Fig. 11

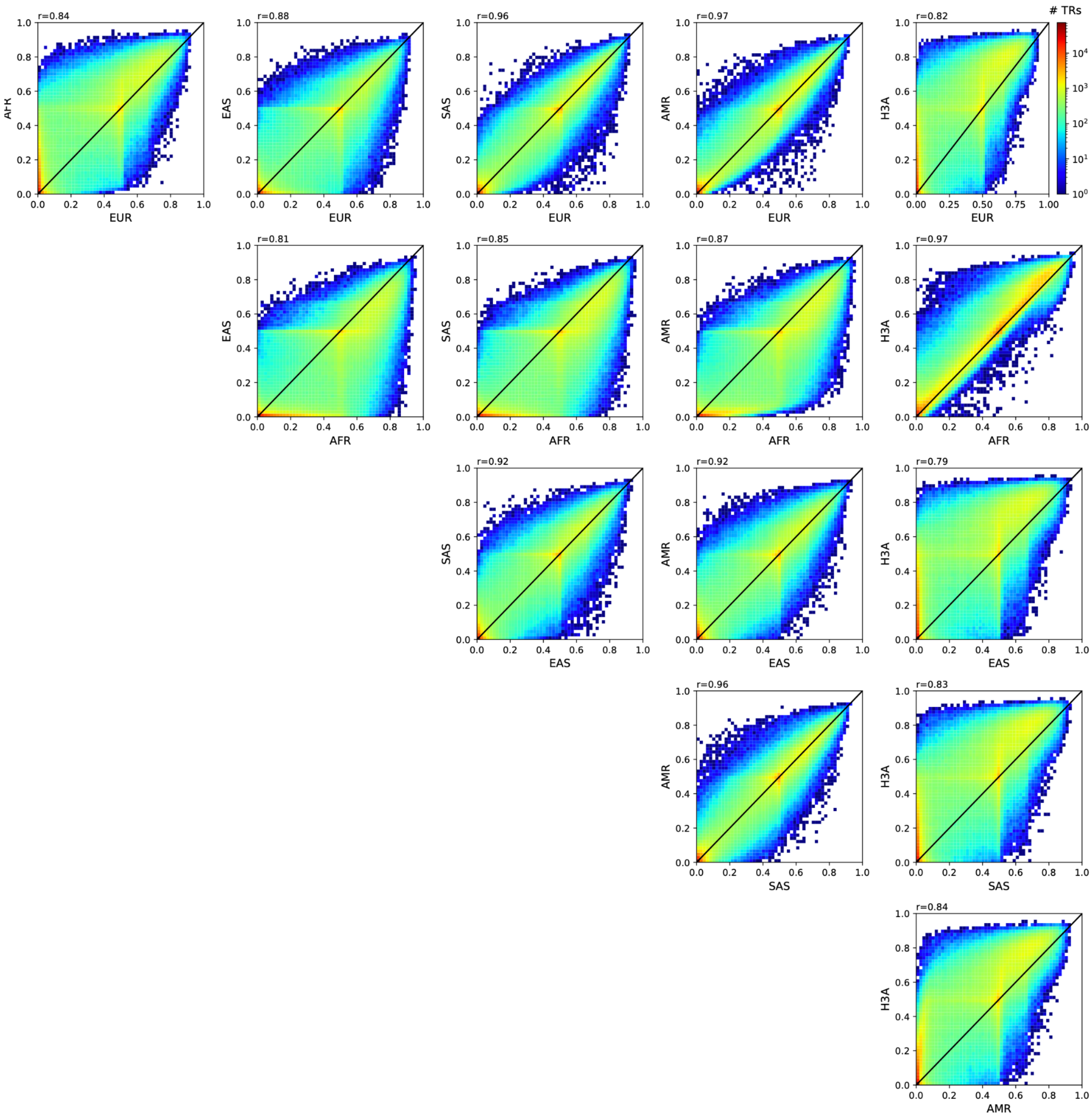


**Comparison of TR heterozygosity values across populations.** Each scatterplot compares the heterozygosity of each TR across two superpopulations. The black line denotes the diagonal. Heatmap shading indicates the number of TRs falling into each bin. The Pearson correlation comparing heterozygosity values in each pair of superpopulations is annotated above each panel. Data is shown for homopolymer TRs.

Supplementary Fig. 12

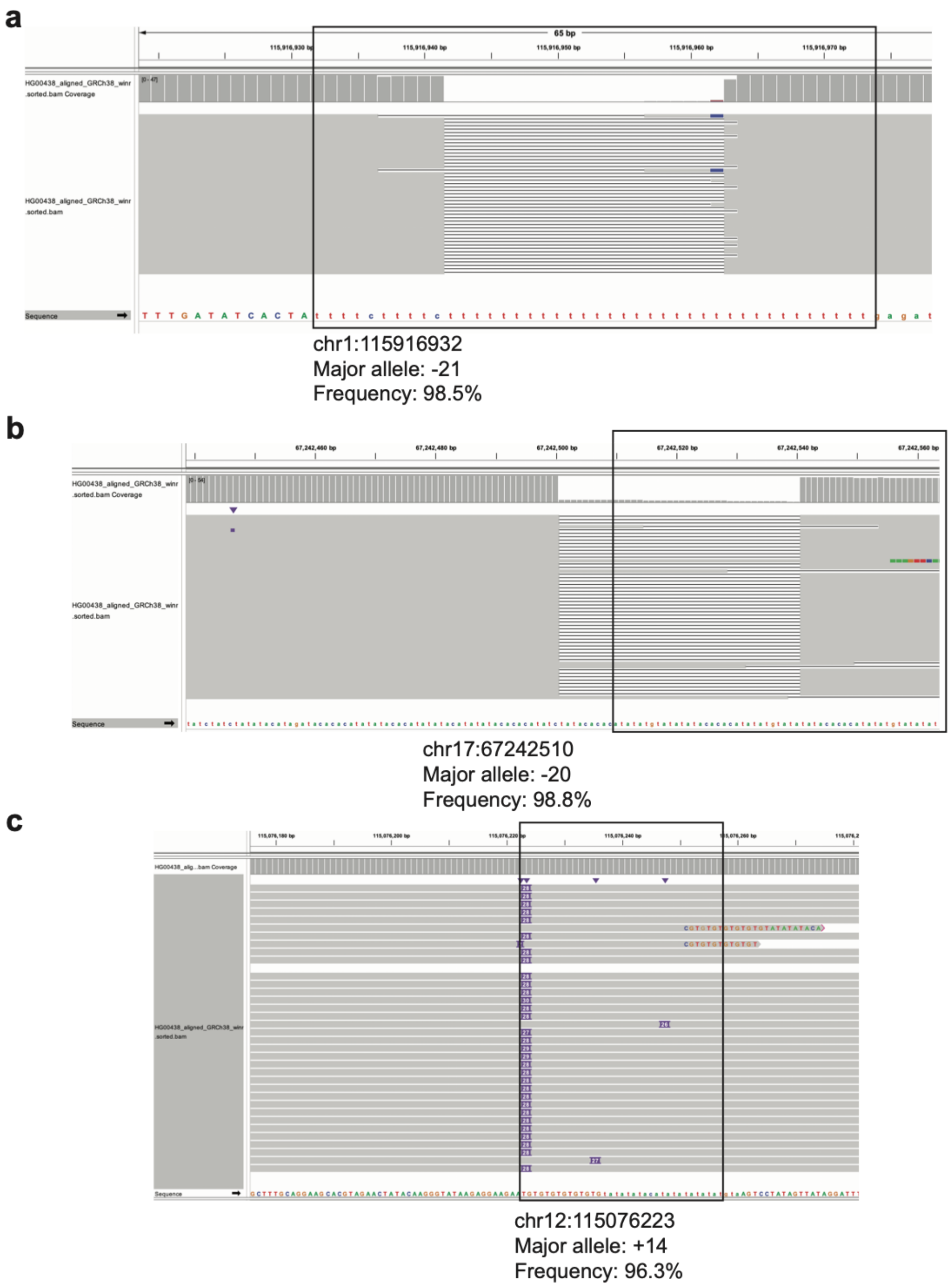


**Example TRs for which the majority of observed alleles show dramatic deviations from the hg38 reference.** In each panel, the TR region is denoted by a black box. Aligned Pacbio HiFi reads from HPRC for sample HG00438 are visualized using the Integrative Genomics Viewer (IGV). The major allele length (repeat units relative to hg38) and frequency are annotated in each panel.

Supplementary Fig. 13

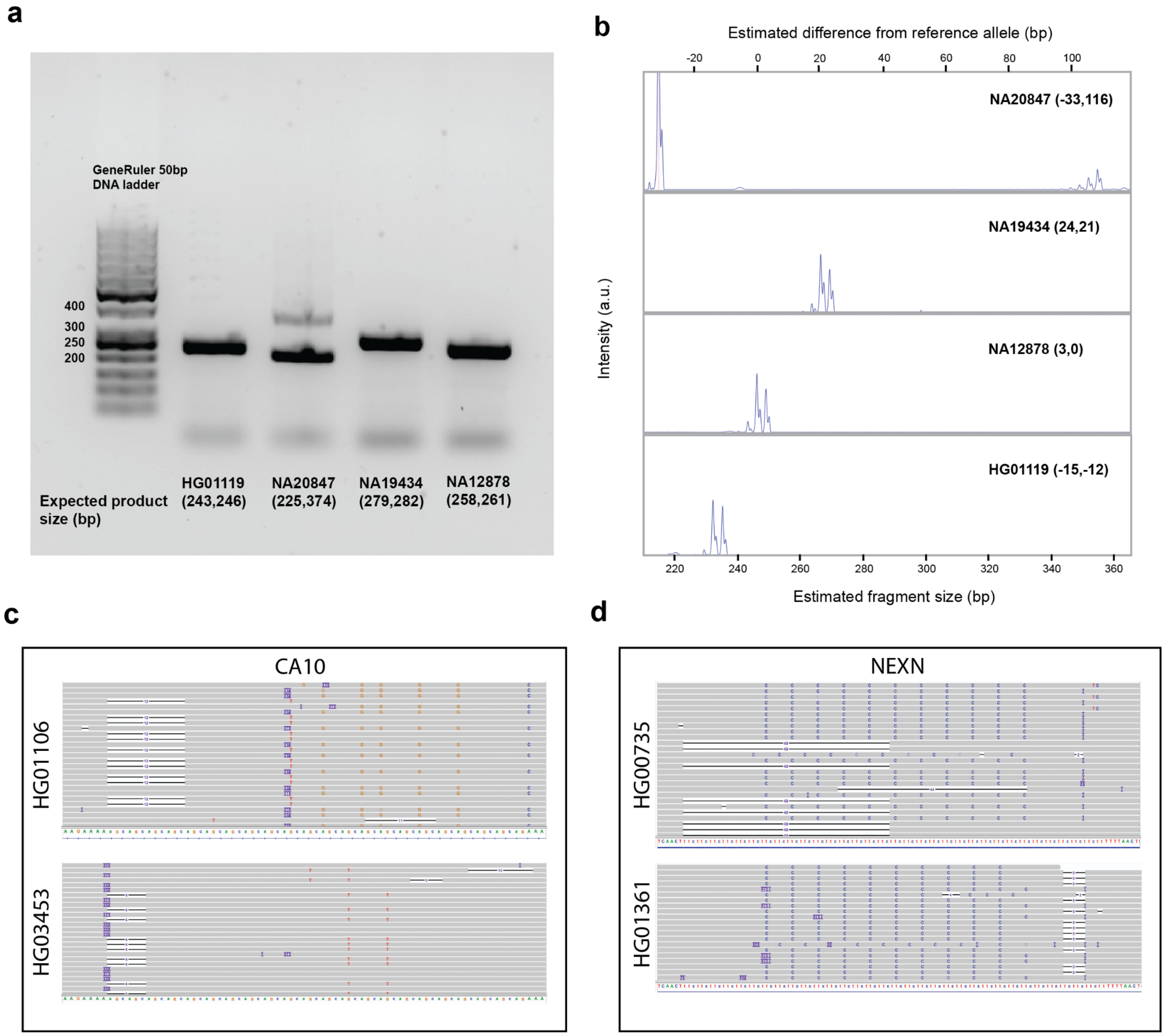


**Experimental validation of genotypes at the CAG repeat in an intron of *CA10*.** In **a.**, PCR was used to amplify the TR region in four 1000GP samples. Sample names and EnsembleTR calls (expected product size in bp) are annotated below each lane. We identified an expansion in NA20847, which is validated by PCR. Panel **b.** Shows results of fragment analysis on the same PCR products using the GeneMapper software. Samples are annotated with EnsembleTR calls (bp difference from hg38). Panels **c-d** show IGV screenshots of Pacbio Hifi reads aligned to the TRs in *CA10* and *NEXN* for which Africa-specific common repeat expansions were identified.

### Supplementary Fig. 14

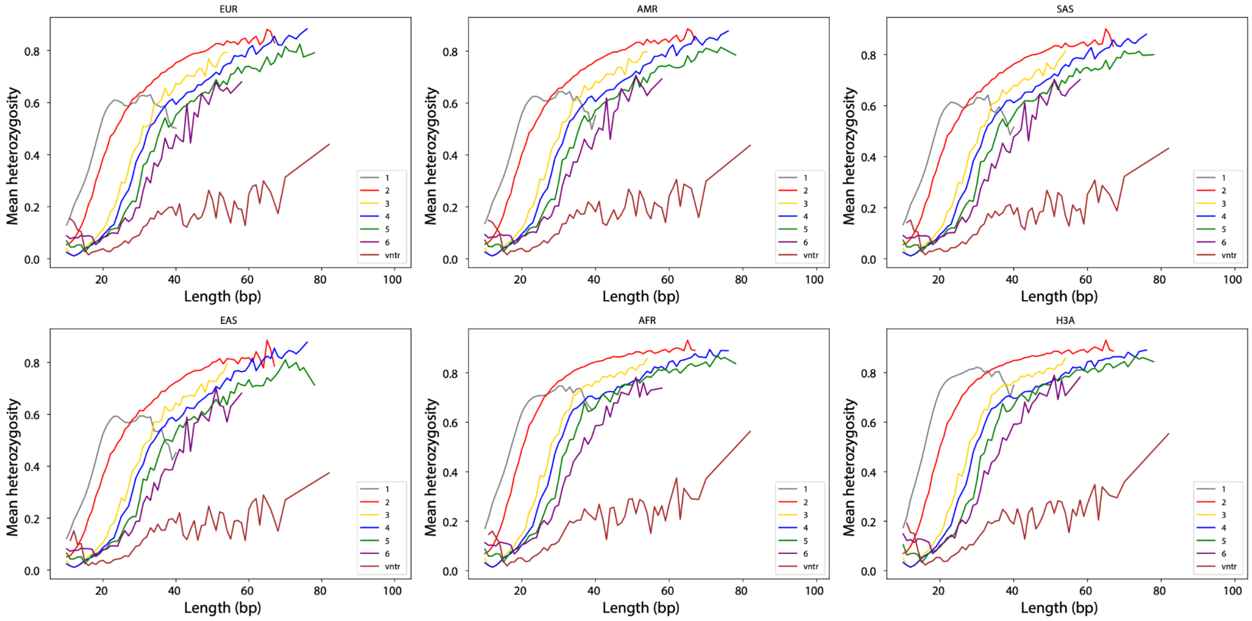


**Heterozygosity is correlated with total TR length.** The x-axis denotes the length of each TR in the reference genome (in bp) and the y-axis shows the mean heterozygosity for TRs with each length. Figures are the same as **Fig. 3a** but plotted separately based on heterozygosity values in each superpopulation. Colors denote different repeat unit lengths. Superpopulation labels are annotated on the top of each panel.

Supplementary Fig. 15

**Stable** / **Polymorphic**

**AC**

Stable:

| Rank | Motif | P-value | log P-pvalue | % of Targets | % of Background |
|---|---|---|---|---|---|
| 1 | TCCTGCT | 1e-19 | -4.394e+01 | 46.41% | 36.25% |
| 2 | GGAGGTGA | 1e-15 | -3.482e+01 | 5.14% | 1.76% |
| 3 | GGCCCTGT | 1e-13 | -3.136e+01 | 19.62% | 13.19% |
| 4 | CATC | 1e-13 | -3.078e+01 | 73.16% | 65.36% |

Polymorphic:

| Rank | Motif | P-value | log P-pvalue | % of Targets | % of Background |
|---|---|---|---|---|---|
| 1 | ACATATATAT | 1e-45 | -1.044e+02 | 28.33% | 15.28% |
| 2 | ACATGCACGCA | 1e-28 | -6.451e+01 | 21.08% | 11.93% |
| 3 | CACATG | 1e-21 | -4.969e+01 | 48.39% | 37.55% |
| 4 | CGCGCAC | 1e-21 | -4.968e+01 | 20.07% | 12.15% |
| 5 | TATACACACAGA | 1e-14 | -3.273e+01 | 6.13% | 2.66% |

**AT**

Stable:

| Rank | Motif | P-value | log P-pvalue | % of Targets | % of Background |
|---|---|---|---|---|---|
| 1 | ATCTTAGCTCAC | 1e-23 | -5.523e+01 | 10.90% | 5.27% |
| 2 | GGGTGCAGTG | 1e-23 | -5.431e+01 | 15.69% | 8.93% |
| 3 | GGCACAG | 1e-23 | -5.372e+01 | 39.62% | 29.78% |
| 4 | GCCGGGCGC | 1e-22 | -5.217e+01 | 12.67% | 6.72% |
| 5 | AGACTCTAT | 1e-22 | -5.180e+01 | 6.78% | 2.60% |
| 6 | TGGCCCAT | 1e-20 | -4.678e+01 | 6.36% | 2.50% |
| 7 | CCTATTTCCC | 1e-18 | -4.190e+01 | 1.15% | 0.00% |
| 8 | TGCTAAGA | 1e-18 | -4.150e+01 | 23.83% | 16.60% |
| 9 | CTTTGAGA | 1e-16 | -3.795e+01 | 19.40% | 13.08% |
| 10 | TGTGCAAATT | 1e-15 | -3.597e+01 | 1.93% | 0.30% |
| 11 | TCACCTCTCC | 1e-15 | -3.590e+01 | 1.15% | 0.03% |
| 12 | AGTGATGGGA | 1e-15 | -3.577e+01 | 4.85% | 1.91% |
| 13 | GCGCT | 1e-15 | -3.560e+01 | 21.53% | 15.12% |
| 14 | ATTTTTCCTATG | 1e-14 | -3.448e+01 | 1.88% | 0.30% |
| 15 | TTCAAGATG | 1e-14 | -3.253e+01 | 2.82% | 0.79% |
| 16 | GCCTGTAAT | 1e-14 | -3.248e+01 | 8.39% | 4.55% |

Polymorphic:

| Rank | Motif | P-value | log P-pvalue | % of Targets | % of Background |
|---|---|---|---|---|---|
| 1 | CACACACACA | 1e-35 | -8.107e+01 | 13.31% | 5.74% |
| 2 | GACATATA | 1e-30 | -6.949e+01 | 22.36% | 13.24% |
| 3 | ACCAGATTT | 1e-17 | -4.068e+01 | 1.58% | 0.05% |
| 4 | TGTAT | 1e-17 | -3.977e+01 | 54.68% | 45.83% |
| 5 | GATTATAAC | 1e-17 | -3.938e+01 | 1.32% | 0.00% |
| 6 | ATACTATTCA | 1e-16 | -3.902e+01 | 2.60% | 0.47% |
| 7 | GGTAGCTTG | 1e-13 | -3.048e+01 | 1.02% | 0.00% |
| 8 | TAATGCCAA | 1e-12 | -2.949e+01 | 0.99% | 0.00% |
| 9 | ATCTATCTATCT | 1e-12 | -2.948e+01 | 1.98% | 0.36% |
| 10 | GAATACCA | 1e-12 | -2.817e+01 | 9.62% | 5.68% |
| 11 | AAGGAAGAC | 1e-12 | -2.773e+01 | 2.01% | 0.42% |

**AG**

Stable: None

Polymorphic:

| Rank | Motif | P-value | log P-pvalue | % of Targets | % of Background |
|---|---|---|---|---|---|
| 1 | CCACTGCATTCC | 1e-13 | -3.206e+01 | 5.61% | 1.44% |

**Significant HOMER results not marked as likely false positives.** Motifs enriched in the context regions for polymorphic vs. stable were identified using HOMER. Results marked as significant by HOMER are shown. Reverse complements were used such that the AC motif category includes information from all AC and GT motif STRs on the forward strand.

Supplementary Fig. 16

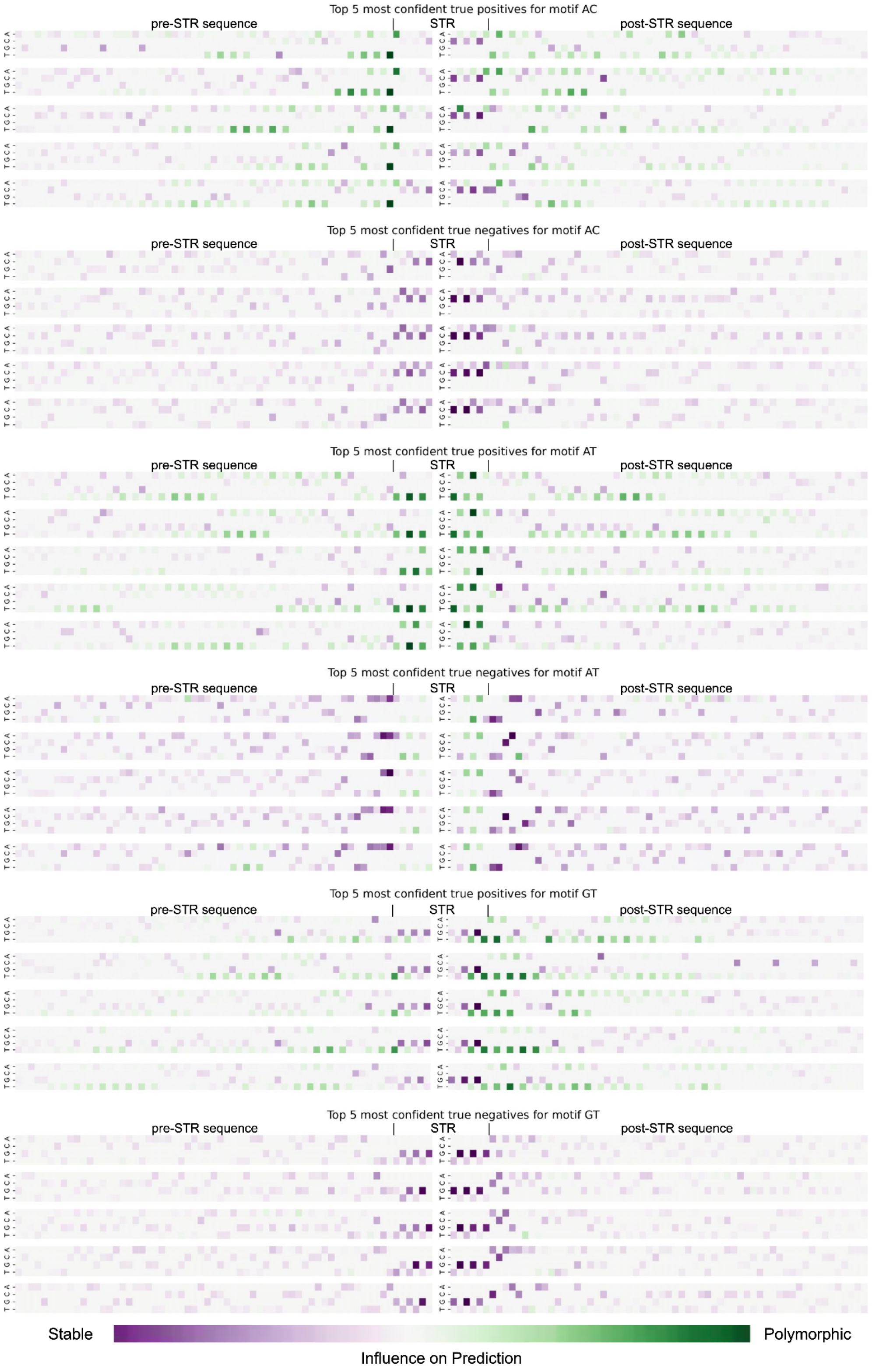

**Integrated Gradients attribution scores for most confidently predicted true positive (correctly predicted polymorphic) and true negative (correctly predicted stable) STRs by the CNN model.** Each row denotes a different TR. Within each row, the matrix has a row for each nucleotide (A, C, G, T) and a column for each position (centered on the TR). Color denotes the attribution score of each base in each position, where green indicates a base positively contributed towards the model predicting polymorphic and purple indicates contributing towards the model predicting stable.

Supplementary Fig. 17

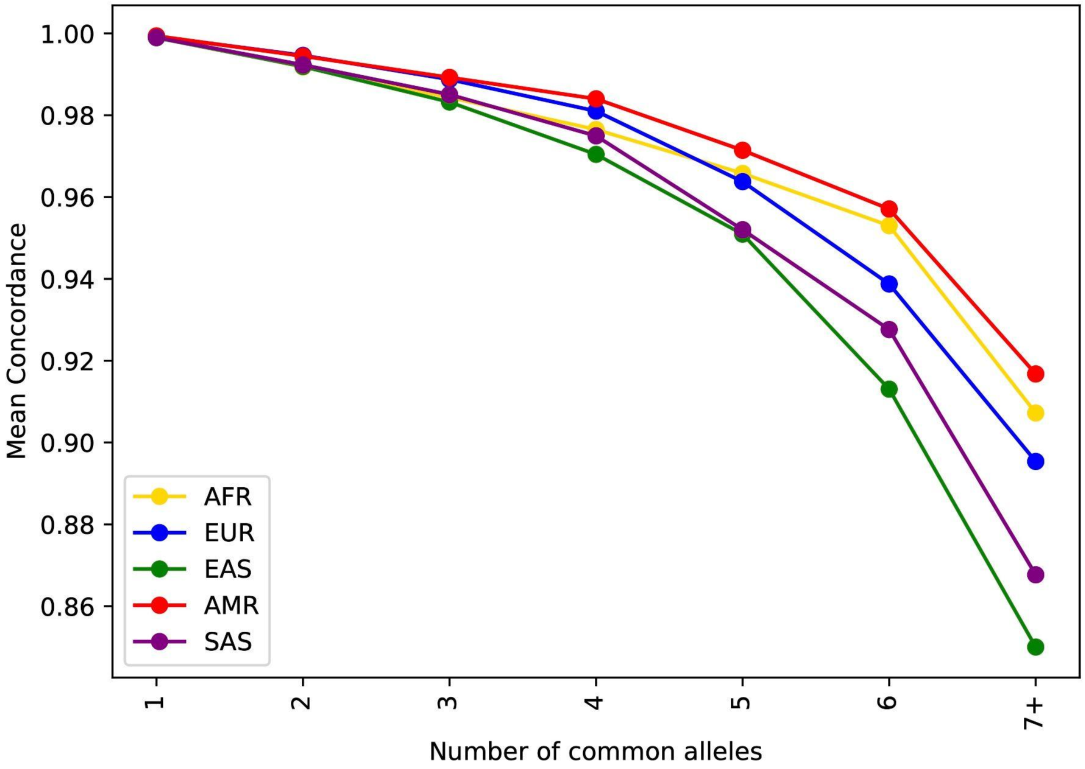


**Imputation accuracy decreases with the number of common alleles.** The x-axis denotes the number of common alleles (frequency > 1%) for each TR. The y-axis denotes the mean concordance for TRs in each bin based on a Leave-One-Out analysis on chromosome 21.
