## Supplementary Dataset 2 for "A deep population reference panel of tandem repeat variation": README

This file has per chromosome tagging SNP information for phased TRs in EnsembleTR callset. 

Phasing TR calls was performed using Beagle v4.0. For each phased TR. The position of SNP with the highest LD with the target TR among nearby SNPs (+-50kb) was reported. 

LD is defined as r^2 where r is the Pearson Correlation Coefficient and it was computed per population (AFR, EAS, SAS, EUR, and AMR). “NA" values mean that all alleles in the corresponding population had the same length. 

Coordinates are based on hg38.

For more information on EnsembleTR please visit its Github Repository:

https://github.com/gymrek-lab/EnsembleTR
